# Base editing derived models of human *WDR34* and *WDR60* disease alleles replicate retrograde IFT and hedgehog signaling defects and suggest disturbed Golgi protein transport

**DOI:** 10.1101/2022.03.14.483768

**Authors:** Dinu Antony, Elif Yýlmaz Güleç, Zeineb Bakey, Isabel Schüle, Gwang-Jin Kim, Ilona Skatulla, Han G. Brunner, Sebastian J. Arnold, Miriam Schmidts

**Author notes:** Correspondence to: Dr. Miriam Schmidts.

## Abstract

Cytoplasmic Dynein-2 or IFT-dynein is the only known retrograde motor for intraflagellar transport, enabling protein trafficking from the ciliary tip to the base. Dysfunction of WDR34 and WDR60, the two intermediate chains of this complex, causes Short Rib Thoracic Dystrophy (SRTD), human skeletal chondrodysplasias with high lethality. Complete loss of function of WDR34 or WDR60 is lethal in vertebrates and individuals with SRTD carry at least one putative hypomorphic missense allele. Gene knockout is therefore not suitable to study the effect of these human missense disease alleles.

Using CRISPR single base editors, we recreated three different patient missense alleles in cilia-APEX-IMCD3 cells. Consistent with previous findings in dynein-2 full loss of function models and patient fibroblasts, mutant cell lines showed hedgehog signaling defects as well as disturbed retrograde IFT. Transcriptomics analysis revealed differentially regulated expression of genes associated with various biological processes, including G-protein-coupled receptor signaling as well extracellular matrix composition, endochondral bone growth and chondrocyte development. Further, we also observed differential regulation of genes associated with Golgi intracellular transport, including downregulation of *Rab6b*, a GTPase involved in Golgi-ER retrograde protein trafficking and interacting with components of cytoplasmic dynein-1, in mutant ciliated and non-ciliated clones compared to controls. In addition to providing cellular model systems enabling investigations of the effect of human SRTD disease alleles, our findings indicate non-ciliary functions for WDR34 and WDR60 in addition to the established roles as components of the retrograde IFT motor complex in cilia.

## Introduction

Dyneins are minus end directed multiprotein motor complexes consisting of heavy-, intermediate-, light intermediate- and light-chains. Powered by ATP, dynein motors transport various cargoes, vesicles and organelles along microtubules. Three different types of dynein motors can be distinguished: axonemal dyneins, dynein-2 (intraflagellar Transport (IFT) dynein) and cytoplasmic dynein (Roberts et al., 2013). While cytoplasmic dynein is essential for retrograde transport along cytosolic microtubules and mitotic spindle arrangement (Canty et al., 2021); axonemal dyneins and IFT dynein are solely found in cilia (Roberts et al., 2013). Axonemal dyneins are essential for ciliary motility and dysfunction of axonemal dynein motor proteins causes primary ciliary dyskinesia (PCD) (MIM #244400), a chronic respiratory characterized by recurrent respiratory infections, laterality defects and sub fertility (Lucas et al., 2020). Dynein-2 is the only known ciliary retrograde motor complex and controls the transport of cargo from the tip of the cilia back to the base (retrograde IFT) (**Figure 1; A**).

**Figure 1.**
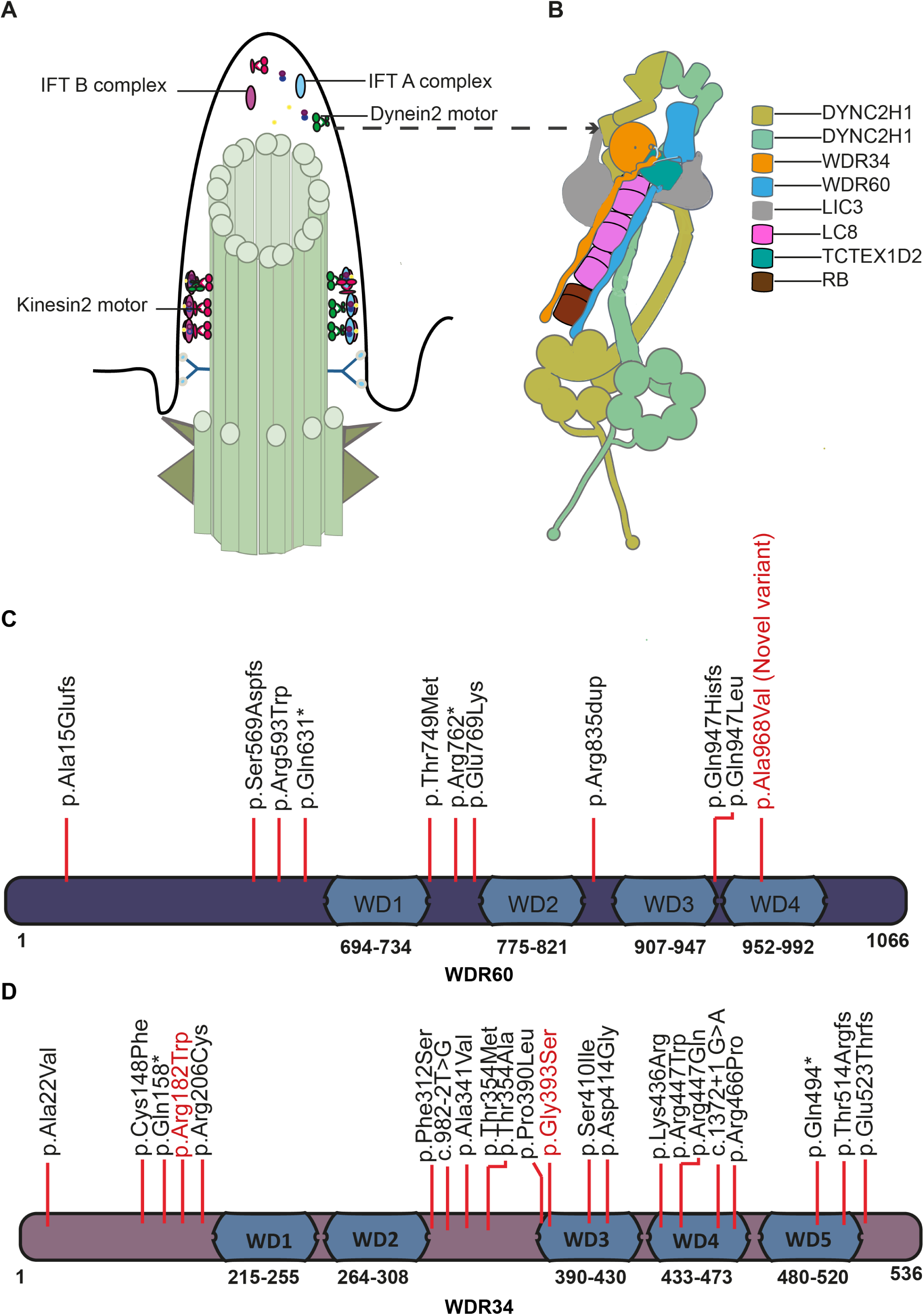
Primary cilia ultrastructure, intraflagellar transport and cytoplasmic dynein-2 composition. and function. **A)** Kinesin-2 motors power the anterograde transport and dynein-2 motors powers the retrograde cargo traffic. IFT-A and IFT-B complexes attach to the motor complexes and serve as cargo adaptors. **B**) Schematic of Dynein-2 motor structure. The two intermediate chains WDR34 and WDR60 bind to the heavy chains as a heterodimeric complex with light chain binding mediated by the N-terminals, adapted from (Antony et al., 2021). **C)** WDR60 protein structure. Known human SRTD disease alleles indicated; the novel not previously described disease variant reported in this paper and recreated in cilia-APEX-IMCD3 cells marked in red. **D**) WDR34 protein structure. Known human SRTD disease alleles; the variants recreated in cilia-APEX-IMCD3 cells are marked in red.

Dynein-2 dysfunction results in a ciliopathy phenotype known as “short rib thoracic dysplasia/dystrophy (SRTD)” in humans. Shortened ribs and long bones resulting in a constricted thoracic cage, pulmonary hypoplasia, and polydactyly are hallmark features of SRTD. Radiologically, so-called handle bar clavicles, a trident appearance of the acetabular roof and cone shaped epiphyses can be observed. Individuals with short-rib polydactyly syndromes (SRPS; MIM #611263, MIM #613091, MIM #263520, MIM #269860, MIM #614091) hereby represent the severe end of the phenotypic spectrum, with affected individuals die prenatally or soon after birth due to cardiorespiratory failure. Jeune syndrome (Asphyxiating Thoracic dystrophy; JATD, MIM 208500) patients present the milder end of the phenotypic spectrum and often survive beyond the neonatal period (Baujat et al., 2013;Schmidts et al., 2013a). While structural heart defects, cystic kidneys and CNS malformations have been described in a subset of SRPS cases, this is not usually the case for JATD individuals (Baujat et al., 2013;Schmidts et al., 2013a).

Recessively-inherited mutations causing SRTD have been identified in most genes known to encode for Dynein-2 subunits including the heavy chain *DYNC2H1*(Dagoneau et al., 2009), the two intermediate chains *WDR34* (Huber et al., 2013;Schmidts et al., 2013b)and *WDR60* (McInerney-Leo et al., 2013), the intermediate light chain *DYNC2LI1* (Taylor et al., 2015) and the light chain *TCTEX1D2* (Schmidts et al., 2015). No mutations have been identified in other light chains to date. A schematic of dynein-2 structure is shown in **Figure 1; B**. *WDR60* variants have been recently reported recently in a family with retinal degeneration and polydactly but without any skeletal dysplasia phenotype (Kakar et al., 2018) while *WDR34* variants have been additionally identified in a case with non-syndromic rod cone dystrophy (RCD) (Solaguren-Beascoa et al., 2021). WDR60 and WDR34 protein structures and localization of published disease-causing variants are shown in **Figure 1; C and D** respectively. Interestingly, all human SRTD cases reported to date carry at least one presumably hypomorphic (mostly missense) allele (Huber et al., 2013;McInerney-Leo et al., 2013;Schmidts et al., 2013b;Zhang et al., 2018). This is in line with the observation that complete knockout of *Dync2h1, Wdr34* or *Wdr60* is lethal around mid-gestation in mice (Ocbina et al., 2011;Wu et al., 2017;Li et al., 2021) and results in severe ciliogenesis defects not observed in patient fibroblasts (Huber et al., 2013;McInerney-Leo et al., 2013).

Hence, to study functional effects of patient alleles, complete gene knockout of WDR34 or WDR60 is unsuitable and recreation of patient missense alleles is needed. We therefore used CRIPSR single base editing to generate mutant cell lines carrying human disease alleles homozygously, including a novel WDR60 disease allele and detected retrograde IFT defects as well as disturbed hedgehog signaling in all mutant clones. GO term analyses of mutant versus control ciliated cells revealed up-regulation of G-protein-coupled receptor signaling as well differentially regulated transcriptions of genes associated with extracellular matrix composition, endochondral bone growth and chondrocyte development. Strikingly, we also observed changes in the expression of a number of Golgi associated genes and found delayed Golgi recovery after monensin mediated stress application in mutants compared to controls. In light with the concordantly observed downregulation of *Rab6b* in all mutant clones, this could indicate that impaired Golgi function and Golgi associated protein transport may contribute to the pathomechanism underlying the patient phenotypes caused by the WDR34 and WDR60 dysfunction.

## Materials and Methods

### Human DNA samples

Written consent was obtained from all participants or their legal guardians. Genetic diagnostic was performed at Radboudumc in Nijmegen under the Diagnostic Innovation Programme Radboudumc Nijmegen. Genomic DNA extraction was performed with Qiagen genomic DNA extraction kit (Qiagen, Germantown, MD, USA). DNA concentrations were determined by Nanodrop (ThermoFischer Scientific, Waltham, MA, USA).

### Whole Exome sequencing

Exome sequencing and data analysis was performed as previously described (Loges et al., 2018;Rad et al., 2019;Rehman et al., 2019). In brief, 2-5 micrograms of DNA from the index case was used for whole exome sequencing (WES) at Novogene, Hongkong. Agilent SureSelect Human All Exon V5 Kit was used for capture and sequencing performed on an Illumina HiSeq 2500 machine. Paired-end sequencing resulted in sequences of 150 bases from each end of the fragments. UCSC hg19 was used as a reference genome. VarScan version 2.2.5 and MuTec and GATK Somatic Indel Detector were used to detect SNV and InDels. Data was then filtered in house using a minor allele frequency (MAF) <1% in public control databases including dbSNP, ExAc and gnomAD remaining variants were first filtered for known disease causing genes first with emphasis on diseases compatible with the patient phenotype (SRTD). Homozygous variants were prioritised due to consanguinity of the family. Additionally, visual BAM file inspection was performed for homozygous CNVs in genes previously associated with SRTD and exome data files were further analysed for presence of heterozygous CNVs using Exome Depth (Plagnol et al., 2012).

### Sanger sequencing

Primers specific for the region were designed using primer-BLAST and PCR amplification of the region was performed with Q5 DNA polymerase (New England Biolabs, Ipswich, Massachusetts, USA). PCR product was then sequenced by reverse primer, using sanger’s chain termination method. Primer sequences are available on request.

### CRISPR/Cas9 mediated mutation creation in Cilia-APEX-IMCD3

Cilia-APEX-IMCD3 (Inner Medullary Collecting Duct) cells were a kind gift from Maxence Nachury, Stanford University School of Medicine, USA. First, we determined corresponding positions of the human disease alleles in the mouse genome (mutation taster programme) (**Figure 3; E-G**). Missense changes created in Cilia-APEX-IMCD3 cell lines are labelled throughout the manuscript with reference to mouse genome. Guide RNA sequences are provided in the **Supplementary table 1**. The selected guide RNAs sequences along with PAM sequence were checked for specificity against the mouse genome (GRCm39) using NCBI BLAST. The gRNA binding regions were checked for presence of single nucleotide polymorphism (SNP) by amplifying the region using One Taq 2X master mix (New England Biolabs, Ipswich, Massachusetts, USA) and subsequent sanger sequencing of the PCR product (primers designed by primer-BLAST, sequences are available on request).The gRNAs along with the overhang sequence were synthesized by Eurofins genomics (Eurofins Genomics, Ebersberg, Germany). The oligos were annealed and ligated into gRNA expression vector, BPK1520 (Plasmid #65777, Addgene, Watertown, Massachusetts, USA) using Anza T4 DNA ligase master mix (Thermo Fisher Scientific, Waltham, Massachusetts, USA) after digesting the vector with BsmBI restriction enzyme (New England Biolabs, Ipswich, Massachusetts, USA).

Cilia-APEX-IMCD3 cells were transfected with 4µg of base editing plasmid BE4-Gam (Plasmid #100806, Addgene, Watertown, Massachusetts, USA) and 1 µg of BPK1520 with respective gRNA insert using jet prime transfection reagent (Polyplus-transfection, Illkirch-Graffenstaden, France) following manufacturer’s instructions. Forty eight hours after the first transfection, cells were split and plated at around 40% confluency and re-transfected the following day. Single cell sorting was performed 72 h after the second transfection (BD FACSAria III Cell Sorter, BD biosciences, Franklin Lakes, New Jersey, U.S.A) and cells were collected in fifteen 96 well plates. For controls, cells were transfected with base editing plasmid BE4-Gam and 1 µg of BPK1520 without any gRNA insert. Single clones were picked approximately 3 weeks later and part of the cells was used for genotyping using Sanger sequencing (primer sequences available on request) (**Figure 2; C**).

**Figure 2.**
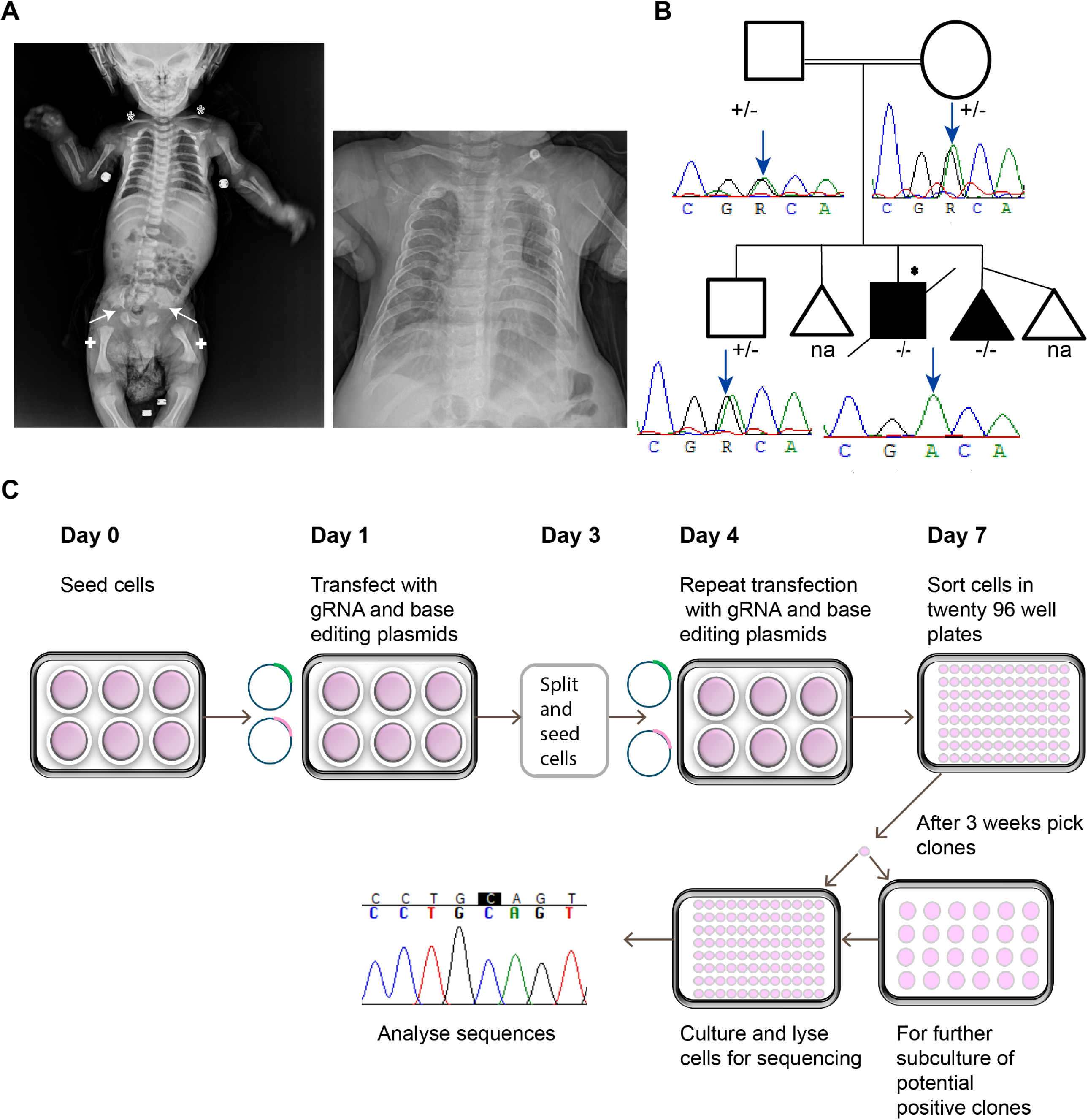
Segregation analysis of the novel WDR60 c. 2903C>T, p.Ala968Val variant identified and CRIPSR single base editing workflow. **A)** Radiographs of the index case showing typical clinical hallmarks of SRTD including shortened ribs, a narrow thorax, shortened long bones (indicated by +), handlebar clavicles (indicated by *) and acetabular spurs (arrows). **B)** Sanger sequencing confirmed that parents and the unaffected sibling carry the allele heterozygously while the affected index (asterix) carries the variant in homozygous state, confirming an autosomal recessive inheritance pattern. **C)** CRIPSR single base editing workflow. Cells were transfected twice with base editing and gRNA expression plasmids. Single cells were FACS sorted 72 hours after the second transfection round and allowed to grow until confluency, followed by clone picking and Sanger sequence analysis to identify edited clones.

### Immunofluorescence analysis

Ciliation was induced by 32 h of serum starvation. Cells were then washed with PBS and fixed with ice cold 100%methanol. Fixed cells were stained with rabbit monoclonal IFT88 (13967-1-AP, Proteintech, Rosemont, IL, USA) and mouse monoclonal gamma tubulin primary antibodies (T6557, Sigma, Taufkirchen, Germany), overnight at 4°C after permeabilizing with 0.5% Triton-X 100. Cells were then washed three times with 1X PBS and incubated with Alexa fluor 568 goat antirabbit (A11011, Life technologies, Carlsbad, CA, USA) and Alexa fluor 633 goat anti-mouse IgG1 (A-21126, Life technologies, Carlsbad, CA, USA) secondary antibodies for 1 h. Cells were finally washed three times with 1 × PBS and mounted in Vecta shield with DAPI (Vector laboratories, Burlingame, USA). Images were taken using Zeiss LSM NLO inverted microscope (Carl Zeiss Microscopy, Jena, Germany) with a 63X objective. Image analysis was performed using Image J software. For investigations of putative retrograde IFT defects, accumulation of IFT88 at the ciliary tip in 130 cilia per mutant clone and control clone was analyzed. For cilia length analyses, the length of the 100 cilia per clone was measured using Image J and for ciliation efficiency 100 cells were checked for cilia formation.

### Western blot analyses

Control and mutant cells were serum starved in medium containing 0.2% FBS for 24 hrs. For GLI3 western blots, cells were then treated with 500nm SAG. After 24hrs, cells were lysed in RIPA lysis buffer (Abcam, Cambridge, UK) followed by total protein concentration estimation (Bradford assay). 30µg of total protein was then separated in NuPAGE Bis Tris gel (4-12%, Invitrogen, Waltham, Massachusetts, USA), followed by western analysis using goat polyclonal GLI3 antibody (AF3690, R&D systems, Minneapolis, Canada). Intensity of bands was then quantified by Image J (densitometry). For PVRL1 western analysis mouse monoclonal antibody was used (MABT61, Sigma, Taufkirchen, Germany) and proteins were separated in Tris Glycine gel 4-20% (Invitrogen, Waltham, Massachusetts, USA). Rabbit polyclonal beta actin antibody was (Ab8227, Abcam, Cambridge, UK) used as loading control in all western analysis.

### RNA sequencing

For RNA sequencing, RNA was extracted from cells grown in 6-well plates using RNeasy Mini Kit (Qiagen, Germantown, MD, USA) following the manufacturer’s instructions. To control for proper ciliation of ciliated samples, a coverslip was placed on one well of the 6-well plate. The cells were then fixed in 4%PFA and stained with rabbit monoclonal ARL13B primary antibody (17711-1-AP, Proteintech, Rosemont, Illinois, USA) and checked for ciliation before RNA extraction. The quality and quantity of total RNA were measured using the High Sensitivity RNA Screen Tape Analysis (Part number 5067-5579, Agilent technologies, Santa Clara, California, United States) and Qubit RNA HS Assay Kit (Thermo Fisher Scientific, Waltham, MA, USA, Q32852) respectively. RNA preparation and RNA sequencing were performed as previously described (Mohammed et al., 2018). RNA sequencing of four different control clones and three different mutant clones with four biological replications each was performed at Novogene, Hongkong on a NovaSeq PE150 sequencer (12 G raw data per sample).

### RNA seq bioinformatics

RNA seq analysis was performed in house as previously described (Tosic et al., 2019). In brief, sequence reads were mapped to the mouse reference genome GRCm38/mm10 (iGenomes, Illumina; chromosomes 1-19, X, Y, M) using the Rsubread v1.28.1 package in R v3.4.4. Gene counts were retrieved from the mapping, applying the Rsubread::featureCounts function with the iGenomes reference genome gtf file for annotation (archive-2015-07-17-32-40 and archive-2015-07-17-33-26 for GRCM38). For differential expression analysis, DESeq2 v1.18.1 was used. First, the counts were normalized by library size (DESeq2::estimateSizeFactors) and by gene-wise dispersion (gene-wise geometric mean over the samples; DESeq2::estimateDispersions), then, the differential expression analysis was performed applying negative binomial generalized linear model fitting and Wald statistics on the normalized count data (DESeq2::DESeq2). Subsequently, the results were filtered for adjusted *p* < 0.05 and the log2 fold change values (log2FC) indicated in the figure legends.

### Golgi stress assay and rescue

Cells were treated with 10µM monensin for 16 h followed by washing twice with 1XPBS and incubating the cells in fresh medium for 6 h. After 6 h the cells were washed in 1XPBS and fixed in 4% PFA followed by staining with rabbit polyclonal beta COP antibody (ab2899, Abcam, Cambridge, UK) after permeabilising with 0.5% Triton-X 100. Alexa fluor 488 goat anti rabbit (A11008, Life technologies, Carlsbad, California, USA) was used as secondary antibody. Cells were imaged and analyzed as explained above. Total 25 cells per condition were counted in one experiment to calculate the dispersed Golgi percentage.

For the rescue experiment mutant cells were first transfected with respective wildtype plasmid. For WDR34p.Arg183Trp and WDR34p.Gly394Ser mutants cells were transfected with WDR34 human Myc-DDK-tagged tagged ORF Clone (RC204288, OriGene, Rockville, Maryland, USA) and for WDR60p.Ala911Val mutant cells were transfected with WDR60 Mouse Myc-DDK-tagged tagged ORF Clone (MR217536, Origene, Maryland, USA). After 24 h cells were treated with 10µM monensin and immunofluorescence analysis was done as explained above. To visualize transfected cells in rescue experiments, mouse monoclonal FLAG primary antibody (F1804-50UG, Sigma, Taufkirchen, Germany) and Alexa fluor 568 Goat anti-mouse secondary antibody (A21144, Life technologies, Carlsbad, California, USA) were used along with COP antibody and corresponding secondary antibody.

## Results

### Identification of a novel disease causing WDR60 allele using Exome Sequencing

Individual SI_36, offspring of consanguineous Turkish parents, was referred to us for genetic analysis from a local genetics clinic, presenting with clinical signs of Short rib thoracic dysplasia (SRTD) including short ribs and long bones, btachydactyly, a narrow thorax, handlebar clavicles, acetabular spines on pelvis x-ray examinations as well as recurrant lower respiratory tract infections and respiratory insufficiency to which he succumbed aged 18 months (**Figure 2A**, a detailed description of the clinical course can be found in the supplementary data file). Exome sequencing revealed a novel homozygous *WDR60* variant (NM_018051.5, c. 2903C>T, p.Ala968Val), not previously described as disease causing in humans and not reported in gnomad, indicating it is a very rare allele. The variant is located at the WD domain of the protein (**Figure 1 C**) and segregated in an autosomal recessive fashion in the family (**Figure 2 B**). The variant is predicted to be disease causing by mutationtaster (64) and probably damaging by polyphen (0.991).

### Recreation of human WDR34 and WDR60 missense alleles using CRISPR base editing

In order to investigate functional effects of the novel *WDR60* allele as well as previously published *WDR34* SRTD patient missense alleles, we proceeded to perform CRISPR base editing (**Figure 2 B**) to recreate human disease causing mutations in Cilia-APEX-IMCD3 cells (Mick et al., 2015). We were able to recreate two *Wdr34* missense alleles homozygously: *Wdr34* NM_001008498.2 (ENSMUST00000113711.3) c.547>T, p.Arg183Trp and c.1180G>A, p.Gly394Ser (in human: WDR34 NM_052844.3:c.544C>T, p.Arg182Trp and c.1177G>A p.Gly393Ser) as well as the novel WDR60 allele we identified in proband SI_36.1 (human c. 2903C>T, p.Ala968Val): mouse *Wdr60* NM_146039.3 (ENSMUST00000039349.8), c.2732C>T p.Ala911Val. When recreating the intended C-T change in *Wdr60*, the cytidine deaminase created an additional heterozygous C-to-T change 7 bases downstream of the targeted base, resulting in a heterozygous synonymous change (c.2739C>T, p.Phe913Phe), hence likely without functional consequences. Likewise, an unintended additional heterozygous change (*Wdr34* c.1180G>A) causing a glycine to aspartic acid change at position 393 was created in the *Wdr34* c.1180G>A, p.Gly394Ser clone. As functional consequences cannot be excluded for this change, we decided to re-create a second missense allele in Wdr34, c.547>T, p.Arg183Trp (**Figure 3 A**). Base editing efficiencies were 0.66% for *Wdr60*, c.2732C>T, p.Ala911Val, 2% for *Wdr34* c.547>T, p.Arg183Trp and 13% for *Wdr34* c.1180G>A, p.Gly394Ser. Protein structure predictions using Phyre2/Missense3D online tools (https://missense3d.bc.i.ac.uk/missense3d/) suggest that the WDR34p.Gly394Ser modification changes the confirmation of the protein due to replacement of a bended buried glycine with an exposed serine, the other two dynein-2 changes did not show any change in the protein conformation. All three changes locate to the WDR34 surface at the interface to DYNC2H1, possibly disturbing protein-protein interactions (**Figure 3 B-D**) and the affected protein positions are evolutionarily well conserved across organisms (**Figure 3 E-G**).

**Figure 3.**
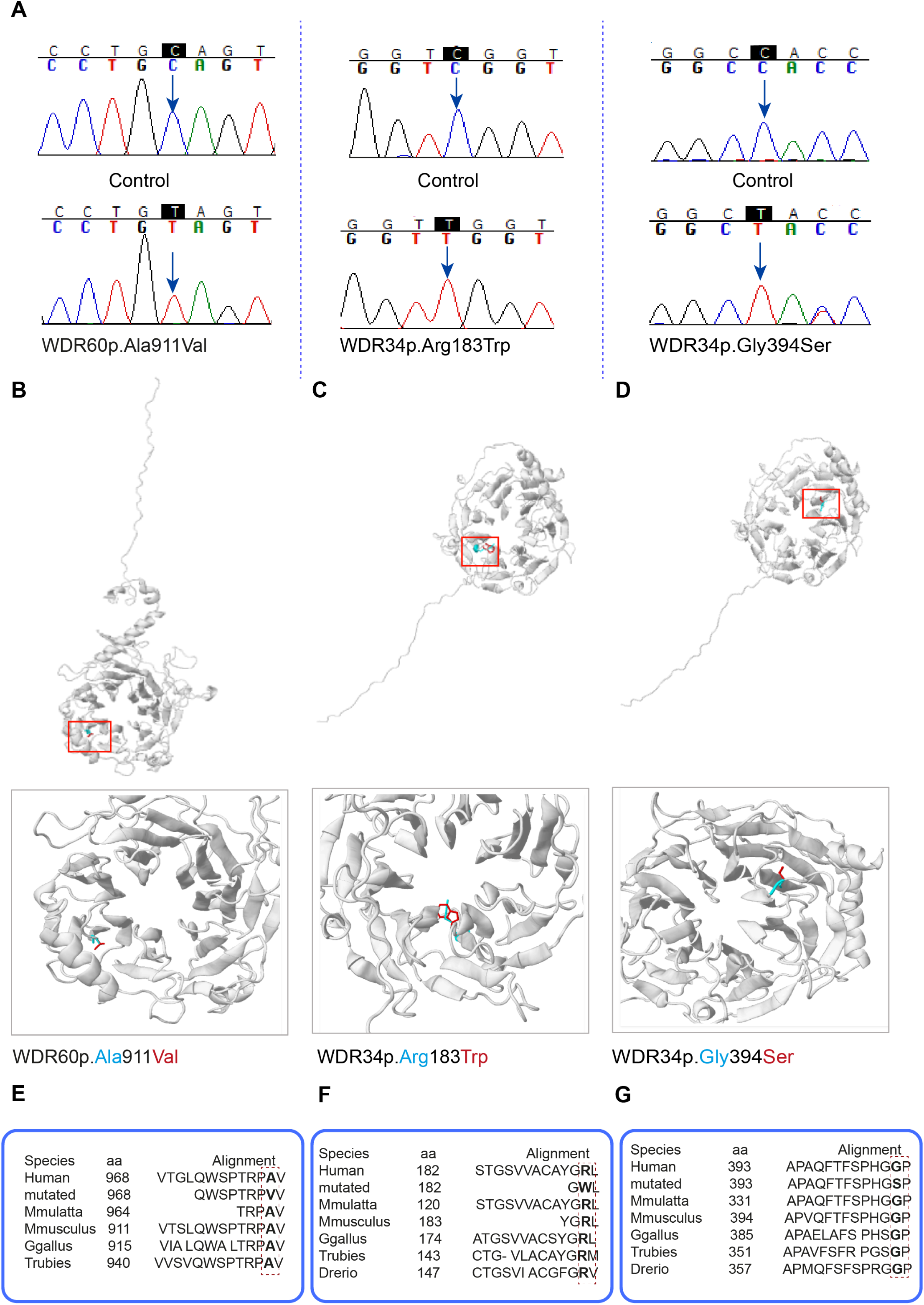
A) Replication of human disease alleles using CRISPR base editing. **A)** Genotypes of homozygously edited clones displaying human disease alleles. Chromatograms depict the single base changes introduced (arrow heads). **B**-**D)**. Protein modelling of generated WDR34 and WDR60 variants. All variants locate to the protein surface. WDR60 p.Ala911Val and WDR34 p.Arg183Trp are predicted to not affect the conformation of the protein, while WDR34p.Gly394Ser is predicted to affect the protein confirmation by exposing the serine residue which replaces a buried glycine. **E-G)**. Cross-species conservation of amino acid positions affected by introduced human disease alleles.

### Retrograde IFT trafficking defects and perturbed hedgehog signaling in Wdr60 and Wdr34 mutant clones

To analyze potential ciliary defects in the three created dynein-2 intermediate chain mutant clones, we proceeded to investigate retrograde IFT visualized by immunofluorescence analysis of the IFT-B component IFT88. Mutant clones showed accumulation of IFT88 at the ciliary tip in more than 50% of cilia compared to 23% of cilia of control clones (**Figure 4 A-E**). Clone WDR34p.Arg183Trp showed somewhat shorter cilia with the majority of cilia in the range of 3µm when compared to controls where most cilia were found to exhibit a length of 4 µm and above. WDR34p.Gly394Ser and WDR60p.Ala911Val ciliary length was similar to controls (**Figure 4 F**). Percentage of ciliated cells was similar amongst mutants and controls (**Figure 4 G**).

**Figure 4.**
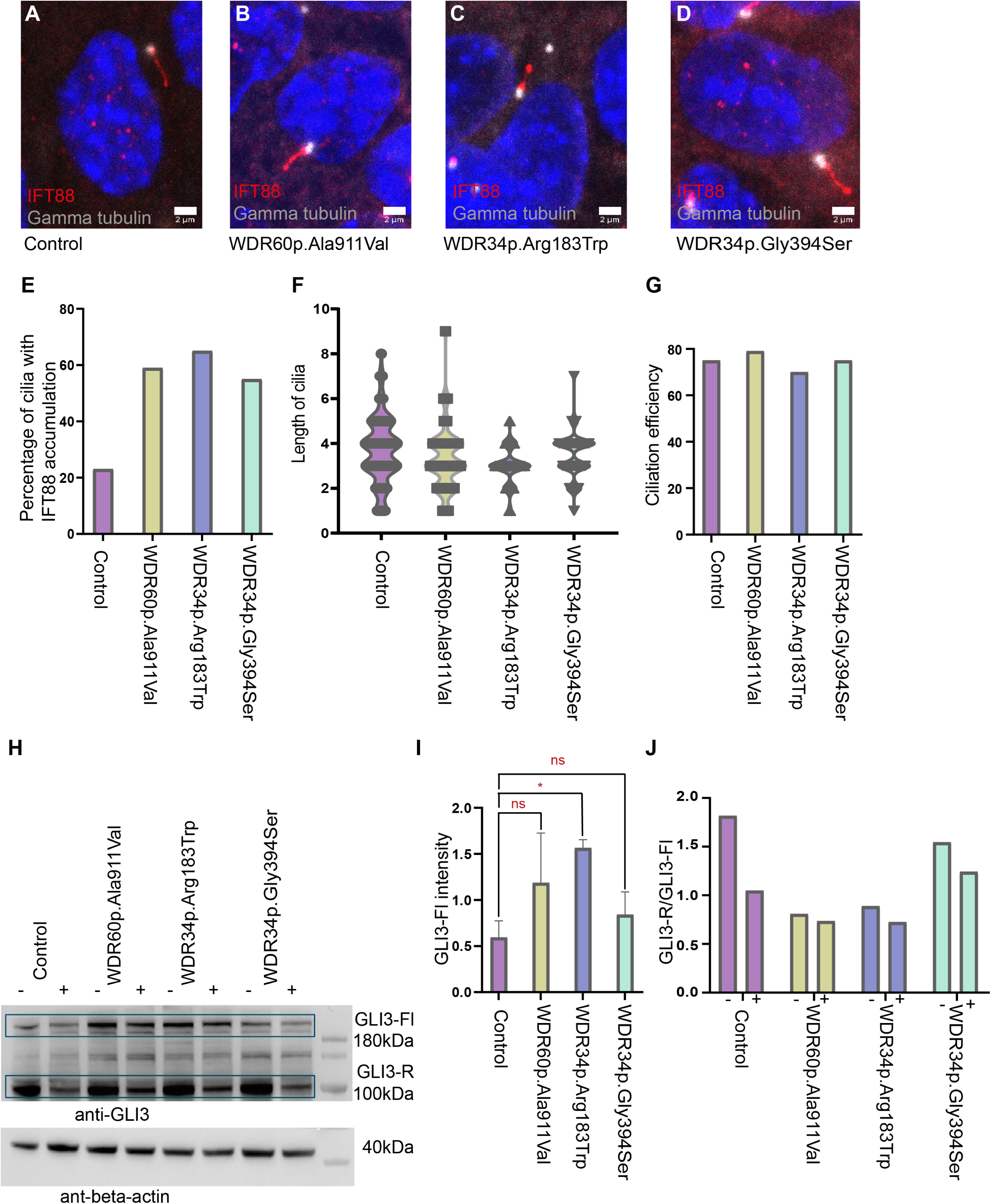
Cellular phenotyping of generated mutants reveals no overt ciliogenesis defects but defective retrograde IFT and disturbed hedgehog signaling. **A-D)** Immunofluorescence analysis of the IFT-B component IFT88 reveals accumulation at the ciliary tip, indicative of defective retrograde IFT. Ciliation was induced by serum starvation. Immunofluorescence analysis, the ciliary base is marked by gamma tubulin (Grey), IFT88 shown in (red). Images were taken using a Zeiss LSM NLO inverted microscope microscope. Scale bars: 2µm. **E)** Quantification of IFT88 accumulation at the ciliary tip in mutant versus control cells. n= 130 cells analyzed per genotype. More than 55% of cilia in dynein-2 intermediate chain mutants display accumulation of IFT88 at the ciliary tip, compared to 23% in controls. **F)** Cilia length distribution in dynein-2 intermediate chain mutants compared to controls. Average length of the cilia in dynein-2 intermediate chain mutants appeared similar to controls, however WDR34p.Arg183Trp cell cilia did not reach maximum lengths observed in the other clones. n=100cells per genotype analyzed **G)** Ciliation efficiency of dynein-2, intermediate chain mutants appeared similar to controls, with at least 75% of ciliation detected in all clones. n=100 cells per genotype analyzed. **H)** Western blot analysis revealed increased GLI3 full length (GLI3-Fl) levels in all dynein-2 intermediate chain mutants compared to controls with and without addition of the hedgehog inductor SAG while baseline GLI3 repressor (GLI3-R) levels appeared unchanged. **I)** Quantification of GLI3-Fl baseline levels in mutants compared to controls. Densitrometry measurements were done using image J. GLI3-Fl levels were further normalised with corresponding loading control. (One way annova with Dunnett’s multiple comparisons test). **J)** Densitomtery analysis of western blot bands and calculation of GLI3 repressor to GLI3 full length ratios amounts before (first bar) and after SAG treatment (second bar) revealed a reduced response in mutant cells compared to controls.

All mutant clones showed perturbed hedgehog signaling in comparison to controls. While application of the hedgehog signaling agonist SAG resulted in a considerable reduction of the GLI3 repressor to GLI3 full length ratio in control cells as expected, this was not observed in WDR34p.Arg183Trp and WDR60p.Ala911Val clones and the response in WDR34p.Gly394Ser cells was markedly reduced when compared to control (**Figure 4 H and J, Supplementary Figure 1**). Further, in line with previously reported findings in *Wdr34* mutant mouse embryos (Wu et al., 2017), full length GLI3 levels were higher in all dynein-2 intermediate chain mutants compared to control cells (**Figure 4 H and I, Supplementary Figure 1)**.

### WDR34 and WDR60 human disease alleles influence gene transcription and suggest dynein-2 intermediate chain dysfunction affects Golgi protein transport

To investigate effects on gene expression resulting from WDR34 and WDR60 patient alleles, we next performed RNA sequencing of the three mutant clones and four different controls. To dissect putative functions related to ciliogenesis, we chose to analyze both ciliated as well as non-ciliated cells. To minimize clonal effects and false positive hits, we investigated all mutants versus all controls for ciliated cells while WDR34p.Gly394Ser was not included in the non-ciliated cells analysis as we performed this run before we obtained the mutant. We observed a higher number of differentially expressed genes between mutants and controls in non-ciliated cells compared to ciliated cells with a greater number of genes downregulated in mutants compared to controls in non-ciliated cells. In contrast, more genes appeared up regulated in mutants vs controls in ciliated cells (**Figure 5 A-C**). Potentially, WDR34p.Gly394Ser not being included in the transcriptomics analysis of non-ciliated cells could contribute towards these differences.

**Figure 5.**
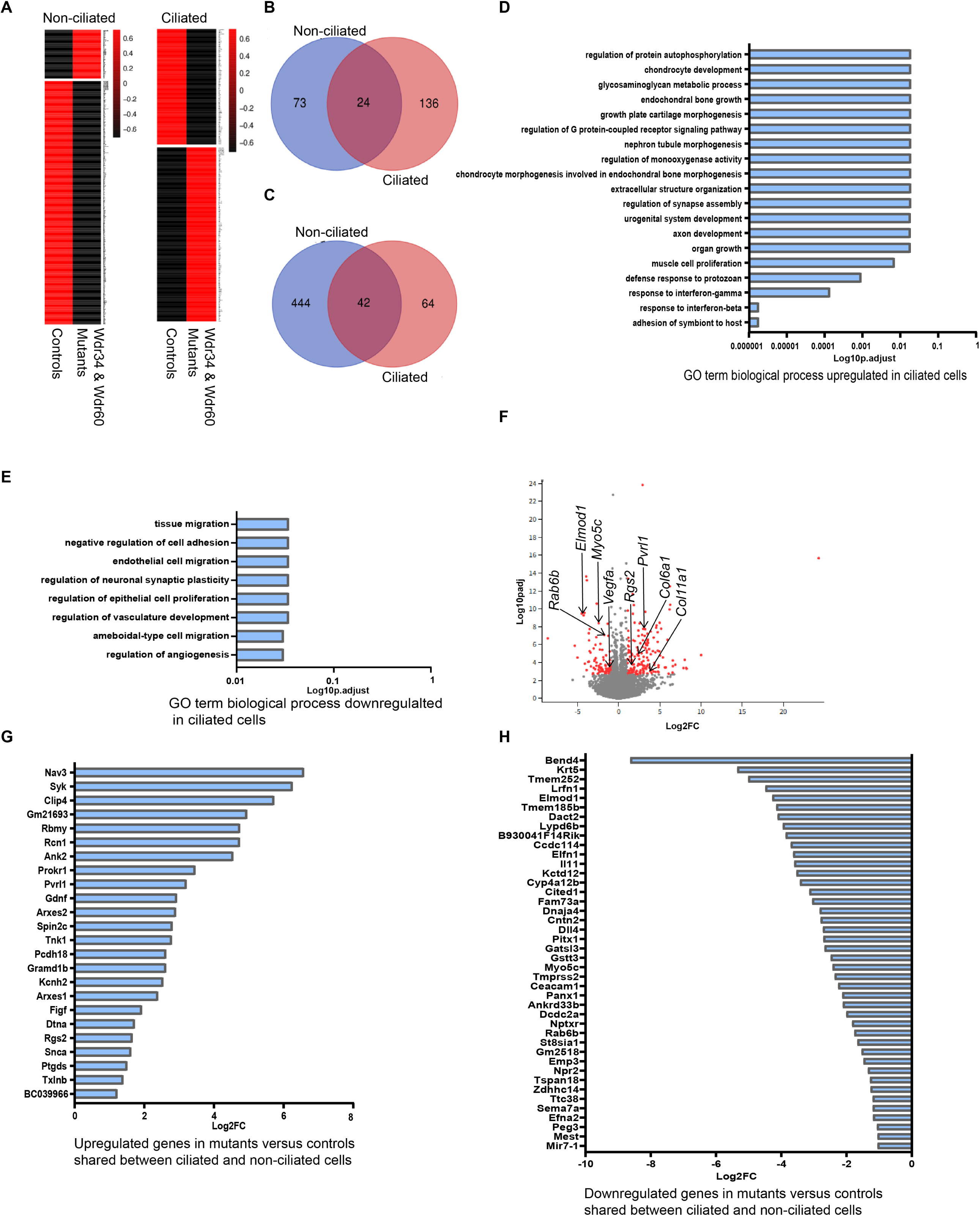
Differential gene expression between dynein-2 intermediate chain mutants and controls. **A)** Heat Map analysis of transcriptome analysis showing a higher number of differentially expressed genes in non-ciliated cells compared to ciliated cells with more genes downregulated in mutants compared to controls. **B**) Venn diagram showing upregulated genes in mutants versus controls of which 24 genes are commonly upregulated in both ciliated and non-ciliated cells. **C)** Venn diagram showing downregulated genes in mutants versus controls with forty-two genes commonly downregulated in both ciliated and non-ciliated cells. **D**) GO term analysis of genes upregulated in mutants versus controls in ciliated cells. **E**) GO term analysis of genes downregulated in mutants versus controls in ciliated cells. **F**) Volcano plot showing differentially expressed genes of ciliated mutants versus controls **G**) Upregulated genes in mutants versus controls shared between ciliated and non-ciliated cells. **H)** Downregulated genes in mutant versus controls shared between ciliated and non-ciliated cells.

We started by analyzing ciliated cells, given the established role of WDR34 and WDR60 for retrograde IFT and cilia function. Gene ontology (GO) term biological process pathway analysis for genes upregulated in mutant ciliated cells compared to controls revealed G-protein-coupled receptor signaling as well as several terms related to skeletal development such as endochondral bone growth, chondrocyte development and extracellular structure organization (**Figure 5D)**. Differentially expressed genes encoding for proteins associated with G-protein related signaling included GTPase activators *Rgs2* and *Rgs9* (De Vries et al., 2000) as well as *Snca*. RGS2 can inhibit COPI (Coat protein I) binding to Golgi and affect Golgi mediated intracellular transport (Sullivan et al., 2000). *Snca* encodes alpha-synuclein, a substrate of G protein-coupled receptor kinases. Synuclein accumulation has been associated with Parkinson’s disease and overexpression of *Snca* in rat ventral midbrain cultures caused reduced neurite length and increased Golgi fragmentation (Furlong et al., 2020). Genes associated with extracellular structure organization, endochondral bone growth and adhesion to symbiont host included *Col6a1, Col6a2, Col11a1, Col16a1* and *Pvrl1* (NECTIN-1). *Pvrl1*, encodes for an actin binding cell adhesion molecule (Barron et al., 2008) (**Supplementary Table 2 and 3**). Western blot analysis confirmed upregulation of PVRL1 in mutant clones compared to controls (**Supplementary Figure 2**).

Downregulated pathways and processes revealed by GO term analysis included angiogenesis/ vasculature development, endothelial cell migration, tissue migration, regulation of epithelial cell proliferation, negative regulation of cell adhesion and regulation of neuronal synaptic plasticity (**Figure 5E)**. Amongst the single genes differentially regulated between ciliated mutants and controls was *Vegfa* (vascular endothelial growth factor-A), playing an important role for angiogenesis (Leung et al., 1989) (**Figure 5E and G)**.

In order to also identify cilia-independent functions of WDR34 and WDR60, we next searched for genes for which expression levels were similarly influenced in ciliated and non-ciliated cells. GO term analysis revealed downregulation of biological processes important for cell and tissue migration (**Supplementary figure 3**). We identified 66 differentially regulated genes between mutants and controls in both non-ciliated and ciliated cells of which 24 were upregulated and 42 were downregulated (**Figure 5; B, C, G and H)**. Amongst those, we observed downregulation of *Dcdc2*, encoding for a protein containing 2 doublecortin peptide domains, localizing to primary cilia and playing a role for tubulin binding, microtubule polymerization and Wnt signaling regulation (Schueler et al., 2015). We further found that several differentially downregulated genes encode for Golgi-associated proteins and proteins involved in vesicle trafficking, including Rab6b (Wanschers et al., 2007), the ARF GAP *Elmod1* (Turn et al., 2021) which has very recently also linked to protein transport from the Golgi to the cilium (Turn et al., 2021), *Myo5c* (Sladewski et al., 2016) and *Zdhhc14* (Sanders et al., 2020) (**Figure 5; G and H**). While GO term analysis did not reveal enrichment of G-protein-coupled receptor signaling, we detected upregulation of two G-protein signaling pathway associated genes *Rgs2* and *Snca*. Likewise, *Pvrl1* was found to be upregulated in both non-ciliated cells as well as ciliated cells while we did not observe an enrichment of extracellular structure or cytoskeleton genes in general.

To further investigate putative Golgi transport defects in dynein-2 intermediate chain mutant cells, we first performed immunofluorescence analyses of COPI vesicles in mutant cells versus controls at basal state. This did not reveal obvious differences (**Figure 6; A-D**). Assuming that mutant cells may exhibit only mild defects due to the hypomorphic nature of the mutations, potentially only detectable under stress conditions, we proceeded to treat the cells with monensin, an agent causing dispersal of the Golgi apparatus and then observed restoration of the Golgi structure after re-application of standard cell culture medium (Oku et al., 2011). This revealed a delayed/impaired Golgi recovery for all three dynein-2 intermediate chain mutants compared to controls (**Figure 6; E-I**). We also observed increased cell death for mutant cells in comparison to controls (data not shown). Overexpression of wildtype WDR34 or WDR60 in the corresponding mutant cell lines partially rescued the delayed/impaired Golgi recovery after monensin treatment (**Figure 6; J-M**).

**Figure 6.**
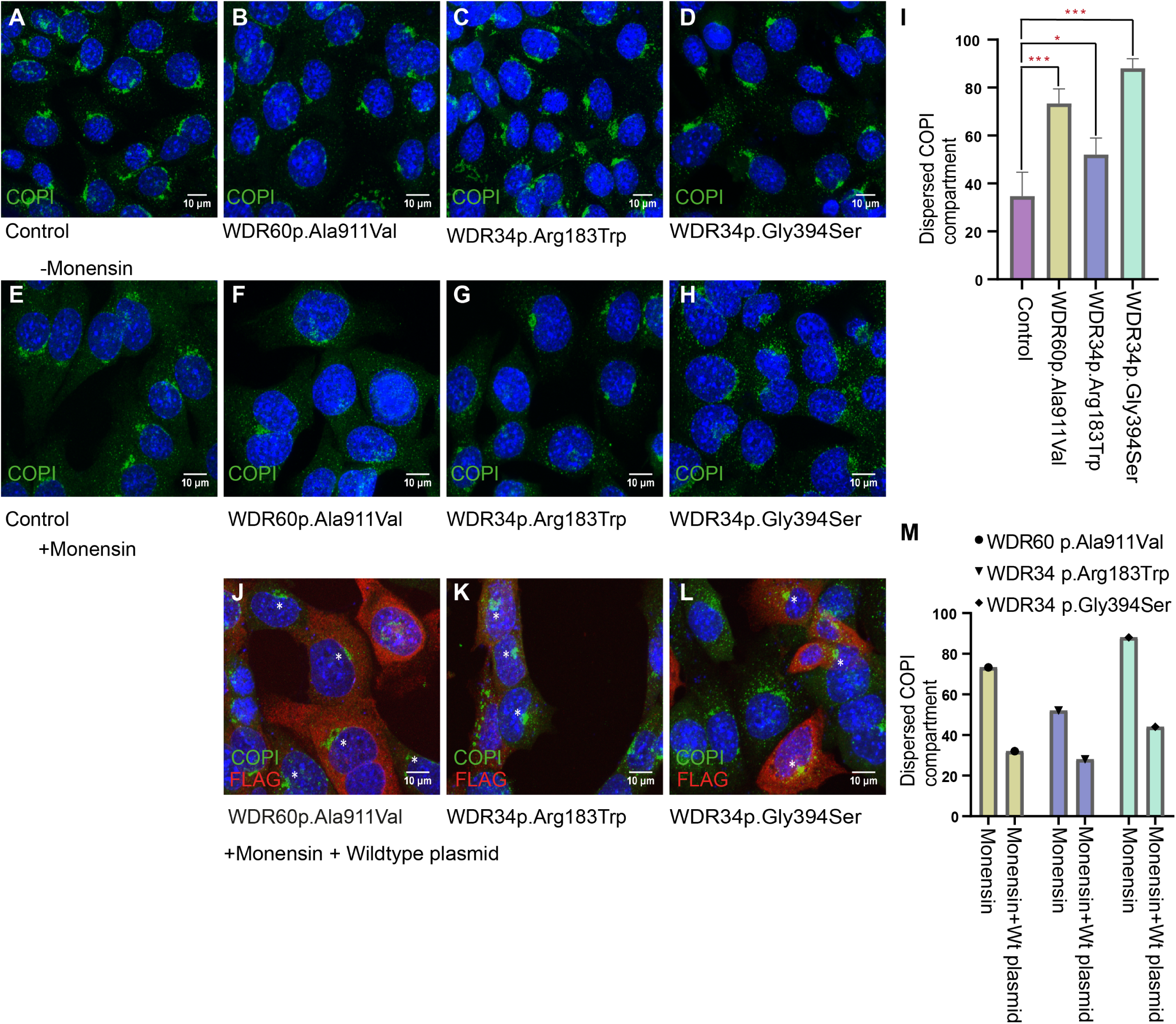
Impaired COPI-Golgi recovery after Golgi stress in mutant versus control clones. **A-D)** COPI immunofluorescence analyses revealing no overt difference between mutant and controls under normal conditions. **E-H)** Monensin treatment followed by washout reveals delayed recovery of the COPI-positive compartment in mutants compared to controls. **I)** Quantification of immunofluorescence results obtained by calculating the ratio of cells with dispersed COPI compartment after the monensin stress assay in mutants versus controls (One way annova with Dunnett’s multiple comparisons test, error bar depicts SD between three independent experiments). **J-M)** Overexpression of Flag-tagged wildtype WDR34 or WDR60 (in red) in corresponding mutant clones partially rescues the dispersed COPI compartment phenotype in the mutants resulting from monensin treatment (asterisks indicate recovered cells expressing the wildtype protein), Scale bars: 10µm in all images **M)** Quantification of cells with dispersed COPI compartment after monensin treatment displaying partial recovery in all three mutants after expression of wildtype protein.

## Discussion

The hypomorphic nature of human disease alleles in dynein-2 components suggests that knockout models may be less suitable for functional studies. Using CRISPR base editing, we were able to replicate three human disease alleles in cilia-Apex-IMCD3 cells. However, editing efficiency using a BE4-Gam cytidine base editor system without GFP for sorting of transfected cells (Komor et al., 2017) was very variable and overall low (0.6-13%). Improved versions of base editing plasmid with GFP tags (Koblan et al., 2018) will likely improve efficiencies. We also experienced unintended editing of bases in close distance to the target base in two instances. This underlies the importance of (if possible) choosing a target base in a genomic area with the least number of possible unintended targets.

In line with previous reports of disturbed retrograde IFT in fibroblasts from human SRTD cases (McInerney-Leo et al., 2013) and knockout mouse models (Wu et al., 2017) we observed disturbed retrograde IFT in all three dynein-2, intermediate chain mutants. We also detected impaired reduction of GLI3 repressor to full length ratio upon addition of the hedgehog agonist SAG as well as increased GLI3 full length expression in all three mutants in comparison to control cells.

Transcriptomics analyses did not reveal any influence of the mutations on the expression level of dynein-2 – complex-or IFT-genes. Expression of *Dcdc2a*, a hepato-renal ciliopathy gene, was significantly lower in all mutants compared to controls. DCDC2A co-localizes with acetylated tubulin along the ciliary axoneme as well as spindle microtubules and is thought to play a role in microtubule polymerization (Massinen et al., 2011). DCDC2A also interacts with DVL1-3, suggesting a role in Wnt signaling regulation (Schueler et al., 2015).In line with this, Schueler et al. suggest that DCDC2A dysfunction results in a ciliogenesis defect in kidney cells with constitutive activation of the canonical Wnt signaling pathway (Schueler et al., 2015). While patients harbouring biallelic DCDC2 mutations do not usually exhibit a skeletal phenotype, nephronophthisis (like) or cystic renal disease as well as rarely cystic-fibrotic hepatic changes can be observed in a subset of SRTD patients, especially in fetuses with an overall severe phenotype. Potentially, reduced DCDC2 expression could contribute to renal and hepatic findings in SRTD.

We also observed consistent downregulation of *Rab6b* in all mutant clones versus controls in both ciliated and non-ciliated cells. RAB6B is a small GTPase important for Golgi to ER transport (Opdam et al., 2000), mediated by COPI vesicles (Hsu et al., 2009). RAB6B interacts with Bicaudal D-1, a binding partner of the dynein-1/dynactin complex (Wanschers et al., 2007)as well as with Roadblock-1 (DYNLRB1) (Wanschers et al., 2008) a light chain component of both cytoplasmic dynein-1 and dynein-2 (IFT-dynein). RAB6B-Bicaudal-D1 complex is proven to be co-localizing at the Golgi (Wanschers et al., 2007). In dynein-2, DYNLRB1 interacts with the N-terminals of WDR34 and WDR60 and this interactions enables the conformational change of dynein-2 motor during IFT transport as well as cargo binding (Asante et al., 2014;Toropova et al., 2019). RAB6B shows 91% similarity with RAB6A and both interact with Rabbkinesin-6 (Opdam et al., 2000) with no specific RAB6B antibody available. Unfortunately, this prevented us from evaluating protein levels and subcellular localization of RAB6B in mutant clones versus controls, however our results suggest impaired Golgi recovery after stress in mutant cells. In line with impaired Golgi protein transport functions under stress, we observed reduced expression of several other Golgi-associated proteins including *Elmod1* in all dynein-2 mutants (both ciliated and non-ciliated cells) compared to controls. The GTPase activating protein ELMOD1 is a GTPAse activating protein which acts mainly on ADP-ribosylation factor (ARF) family GTPases and knock out affects primary cilia formation, reducing ciliary ARL13B expression and resulting in accumulation of INPP5E and IFT140 at the Golgi (Turn et al., 2021). Knockout mice present with fused and elongated stereocilia in the inner ear and disturbed intracellular vesicle trafficking in knockout cells has been described (Krey et al., 2018). Further, we detected up-regulation of RGS2, a regulator of G protein signaling and inhibitor of COPI vesicles binding to Golgi in all mutants (Sullivan et al., 2000).

Due to the extensive amount of extracellular matrix proteins required for cartilage and bone formation and maintenance, proper function of protein transport and posttranslational modification at the Golgi are essential for skeletal development. It comes as no surprise that defective protein processing and transport as well as Golgi dysfunction has been observed with a number of skeletal dysplasias/developmental skeletal conditions, including geroderma osteodysplasticum (OMIM: 231070), caused by dysfunction of GORAB, a trans-golgi and centrosome RAB6-interacting protein (Egerer et al., 2015) and *ARCN1-* associated syndromic rhizomelic short statur*e* (Izumi et al., 2016). *ARCN1* encodes delta-COP, localizing at the Golgi, ER and intracellular vesicles and plays a role in ER stress regulation and ER-Golgi protein transport.

Previous studies have further suggested links between ciliary proteins, Golgi function and skeletal dysplasias: Grissom et al. have suggested that the dynein-2 light intermediate chain DYNC2LI1 co-localizes with DYNC2H1 at Golgi and mediates cargo-loading to the dynein-2 complex at the Golgi (Grissom et al., 2002). Follit et al. demonstrated that TRIP11 (GMAP-210), a Golgi associated microtubule binding protein, anchors IFT20 to the Golgi (Follit et al., 2008). TRIP11 is able to recruit gamma-tubulin containing protein complexes to the Golgi membrane independent of microtubule integrity and TRIP11 depletion causes Golgi fragmentation (Rios et al., 2004). Dysfunction of TRIP11 causes a severe chondrodysplasia phenotype presenting with thoracic dysplasia resulting in lung hypoplasia in humans and mice (Smits et al., 2010;Wehrle et al., 2019) with cells showing a defective Golgi architecture defective Golgi-mediated glycosylation and intracellular accumulation of the extracellular matrix protein perlecan (Smits et al., 2010).

In line with a mildly perturbed COPI-Golgi function in our mutants, we found reduced COPI-Golgi recovery abilities after monensin stress application in mutants versus controls, that could be partially rescued by expression of wildtype WDR34 or WDR60. Overall, Wdr34p.Gly394Ser mutant cells displayed a more severe phenotype compared to the other two mutants, while described human phenotypes for the two different WDR34 alleles does not seem to differ with regards to severity with death in utero or within the first months of life (Schmidts et al., 2013b;Zhang et al., 2018). However, protein structure modelling using Phyre2 predicted that the WDR34p.Gly394Ser change disrupts the protein 3D structure due to replacement of a bended buried glycine with exposed serine while the other two disease associated mutations are predicted to have less severe functional effect (**Figure 3D**). Potentially, this predicted more disrupting effect of WDR34p.Gly394Ser could account for the more pronounced in vitro phenotype we observed. However, we also cannot exclude additional effects of the unintentional heterozygous p.Gly393Asp change in our clone.

While GO term analyses of mutant versus wildtype clones revealed down-regulation of pathways essential for angiogenesis/vasculature development as well as cell and tissue migration in both ciliated and non-ciliated cells, we observed lower expression of *Vegfa* only in in mutant ciliated cells compared to ciliated controls. While *Vegfa* knockout is embryonically lethal in mice, hypomorphic loss of function results in angiogenesis defects including cartilage vascularization defects (Zelzer et al., 2002). Reduced *Vegfa* expression has further been found at the the growth plates of *Kif3a* knockout mice. Kif3 encodes for a subunit of the anterograde IFT-motor kinesin and Kif3a loss of function phenotypes in mice ressemble dynein-2 dysfunction phenotypes including hedgehog signaling defects, growth plate dysfunction and skeletal defects (Koyama et al., 2007). This could indicate a role of Cilia or IFT to regulate Vegfa expression.

In summary, in this study, we successfully created SRTD disease models carrying human missense alleles using base editing. We confirmed retrograde IFT defects as well as disturbed hedgehog signaling and further detected evidence for COPI-Golgi perturbation associated with human disease alleles which could, combined with changes in the cytoskeleton and extracellular matrix composition, contribute towards the severe skeletal phenotype associated with dynein-2 intermediate chain dysfunction.

## Web Resources

https://benchling.com/pub/liu-base-editor

https://blast.ncbi.nlm.nih.gov/Blast.cgi

https://www.mutationtaster.org/

https://www.ncbi.nlm.nih.gov/tools/primer-blast/

http://genetics.bwh.harvard.edu/pph2/

http://missense3d.bc.ic.ac.uk/~missense3d/.

## Acknowledgements

We would like to thank the index individual and his family for participation in the study.

