## Supplemental data for "Base editing derived models of human *WDR34* and *WDR60* disease alleles replicate retrograde IFT and hedgehog signaling defects and suggest disturbed Golgi protein transport"

**Supplementary Data**

**Detailed description of the clinical phenotype of SI_36**

The index was the second child of consanguineous couple (second degree cousins). His mother previously had given birth to a healthy boy and reported one miscarriage where no genetic testing was performed. The mother was referred to our clinic for genetic counseling when she was 26week pregnant for the index. On detailed prenatal ultrasonography, the fetus presented with short and mildly bowed extremities (femur, humerus tibia) and mild angulation on humerus, normal head circumference, frontal bossing, mildly narrow thorax (increased cardiothoracic index: 66%) and right pes equinovarus. Chromosome analysis and FGFR3 gene analysis from cord blood were normal. He was born at term with a birth weight of 3400 gr, bt ht: 51 cm, bt OCD: 38 cm and stayed at the NICU for 2 weeks because of respiratory distress and direct hyperbilirubinemia (cholestasis). He presented with a narrow thoracic cage, predominantly rhizomelic short extremities short hands and feet and brachydactyly, partial cutaneous syndactyly of 2nd and 3rd toes, bilateral palmar simian creases, distally placed small nails, especially toe nails were deeply placed and hypoplastic, high palate with normal frenula, mild umbilical hernia and normal genitalia. Measurements at 17 days of age were Wt: 3260 gr, ht: 51.5 cm, OFD: 35.5 cm. He was clinically diagnosed with Short Rib Thoracic Dysplasia /Jeune Syndrome.

Newborn Echocardiography showed mild peripheral pulmonary stenosis, a control at 1 year of age showed no abnormalities. Newborn abdominal ultrasound showed mild hepatomegaly, grade hydronephrosis of the left kidney but no bile duct abnormality. A control showed normalisation of the left sided hydronephrosis. Opthalmological examination and brain ultrasound revealed no abnormalities.

During follow up, he developed several lower airway infections and he suffered from chronic respiratory distress. Sternotomy and thoracic expansion surgery were performed at the age of 7 months. He was last evaluated when he was 14 months old, measurements were height: 69-70 cm (< 3. Centile), weight: 6730 gr (< 3rd centile), OFD: 43.5 cm, when he still experienced respiratory distress, mild cyanosis, narrow thoracic cage with distended abdomen. His nasal base appeared flat, had frontal bossing, hypertelorism, and defective wedged enamels of his teeth. He was admitted several times to the ICU for respiratory insufficiency. A thoracic CT scan at the age of 6 months revealed a narrow thoracic cage especially on superior and anterior parts, relative cardiomegaly, right aberrant subclavian artery, ground-glass opacifications in some segments of both lungs and linear opacities showing linear subpleural atelectasis.

He further suffered from chronic hepatic disease and portal hypertension and he had paracentesis twice for ascites. Portal vein Doppler and ultrasound at the age of 9 months revealed hepatosplenomegaly, parenchymal heterogeneity of the liver, decreased caliber of the portal vein, increased echogenity of periportal regions. Decreased blood flow through the portal vein and ascites.

He was trying to stand up and walk at 14 months of age. His voice was low pitched due to long term intubation. At 1.5 years of age, he sadly passed away.

Three years later the mother was pregnant again with twins, however both fetuses died in utero round five months into the pregnancy. One fetus was macerated at birth. The other fetus showed mildly narrow thorax, short extremities and brachydactyly of the hands. No autopsy was performed.

| |
|---|
| **WDR60 p.Ala911Val**<br>Guide Forward 5’CCTGCAGTGTTCCTGGTCC 3’<br>Guide Reverse 5’GGACCAGGAACACTGCAGG 3’ |
| **WDR34 p.Arg183Trp**<br>Guide Forward 5’ TGGTCGGTGAGTGAGAGC 3’<br>Guide Reverse 5’ GCTCTCACTCACCGACCA 3’ |
| **WDR34 p.Gly394Ser**<br>Guide Forward 5´GGGCCACCATGGGGAGAGA3´<br>Guide Reverse 5´TCTCTCCCCATGGTGGCCC 3´ |

**Supplementary Table 1**. gRNA sequences used for CRISPR single base editing

| **Description** | **geneID** |
|---|---|
| adhesion of symbiont to host | Ace2/Gbp7/Gbp3/Nectin1/Gbp2/Gbp6 |

| | |
|---|---|
| response to interferon-beta | Ifi203/Tgtp1/Tgtp2/Gbp3/Gbp2/Gbp6/Ifi47/Ifitm3 |
| response to interferon-gamma | Tgtp1/Gbp7/Gbp3/Gbp10/Gbp2/Gbp6/Snca/Ifitm3/Gbp4 |
| defense response to protozoan | Gbp7/Gbp3/Gbp10/Gbp2/Gbp6 |
| muscle cell proliferation | Il15/Ace2/Angpt1/Esr1/Tenm4/Retn/Tgm2/Gper1/Fgf1 |
| organ growth | Col6a2/Ddr2/Col6a1/Esr1/Tenm4/Rgs2/Matn2/Fgf1 |
| axon development | Dclk1/Tspan2/Epha7/Map2/Nectin1/Gdnf/Ispd/Flrt2/Adcy1/Ust/Matn2/Bex1 |
| urogenital system development | Angpt1/Kif26b/Epha7/Nfia/Gdnf/Esr1/Serpinb5/Hey1/Irx1/Fgf1 |
| regulation of synapse assembly | Ptprd/Epha7/Adgrl3/Nectin1/Flrt2/Snca |
| extracellular structure organization | Lox/Abca1/Col11a1/Ddr2/Eln/Abi3bp/Serpinb5/Flrt2/Col16a1 |
| chondrocyte morphogenesis involved in endochondral bone morphogenesis | Col6a2/Col6a1/Matn2 |
| regulation of monooxygenase activity | Gdnf/Esr1/Atp2b4/Snca |
| nephron tubule morphogenesis | Kif26b/Gdnf/Hey1/Irx1/Fgf1 |
| regulation of G protein-coupled receptor signaling pathway | Rgs9/Kctd12b/Rgs2/Atp2b4/Snca/Gper1 |
| growth plate cartilage morphogenesis | Col6a2/Col6a1/Matn2 |
| endochondral bone growth | Col6a2/Ddr2/Col6a1/Matn2 |
| glycosaminoglycan metabolic process | Dse/Il15/Hexa/Dcn/Angpt1 |
| chondrocyte development | Col11a1/Col6a2/Col6a1/Matn2 |
| regulation of protein autophosphorylation | Gpnmb/Vegfc/Syk/Epha7 |

**Supplementary Table 2**. Gene ontology term biological process; upregulated in ciliated cells

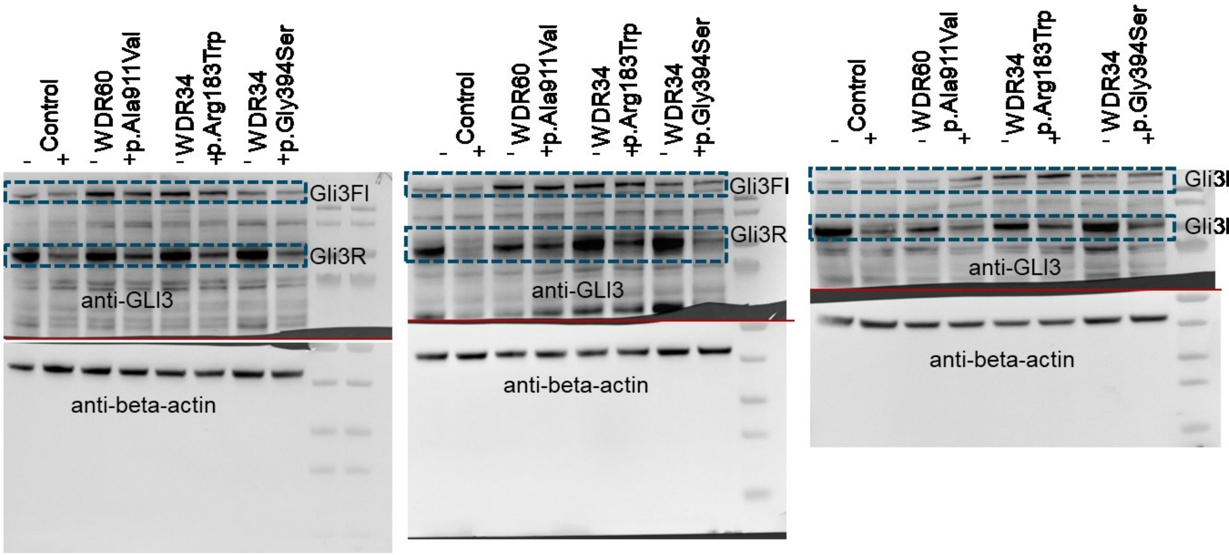


**Supplementary Figure 1. Full gel images showing expression levels of GLI3 full length.** Gli3 repressor and beta-actin as a loading control. Gli3FL: Gli3 full length; Gli3R: Gli3 repressor. n=3 independent experiments. beta-actin was assessed on a separate gel (red line indicates the two separate gels run) due to overlapping size with Gli3.

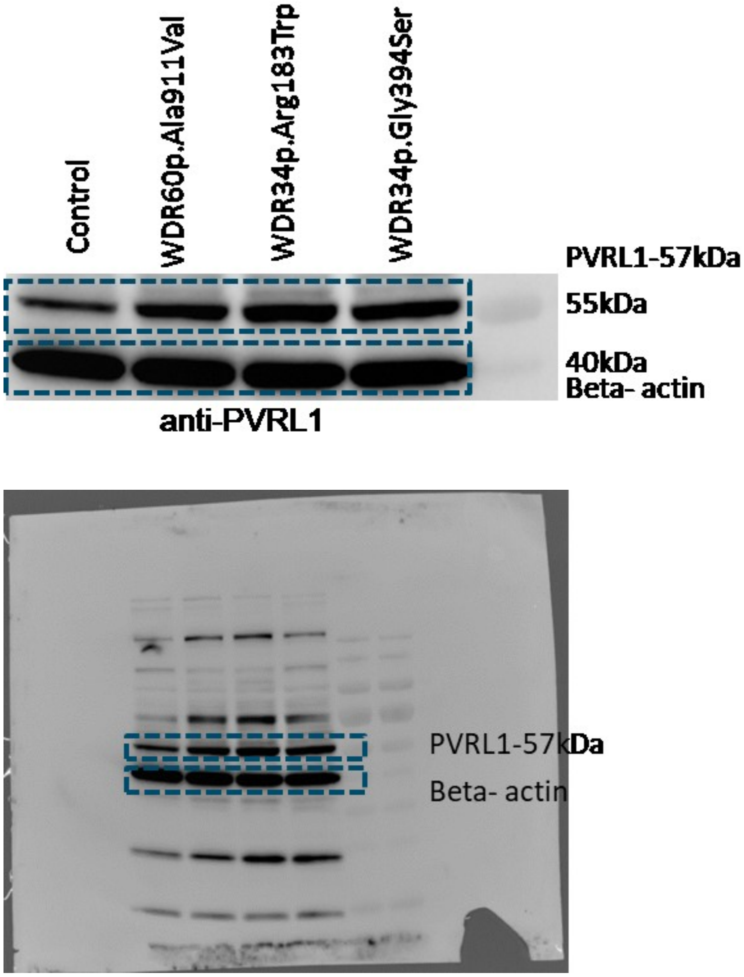


**Supplementary Figure 2. Full gel images of PVRL1 western blot**. Beta actin was used as loading control.

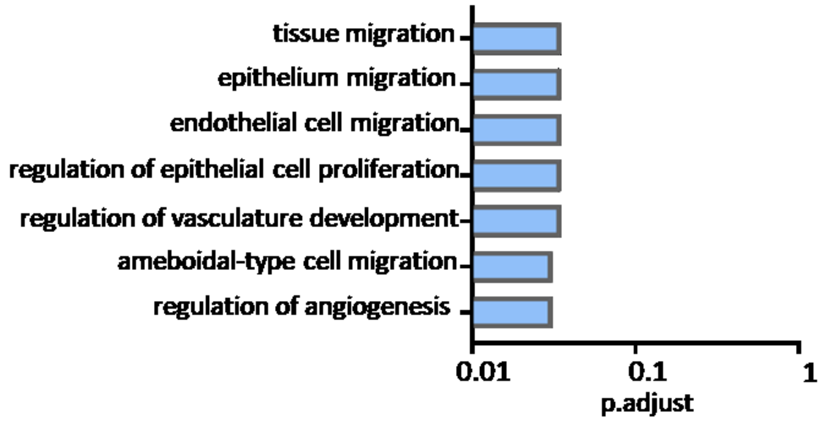


**Figure 3. GO term biological process downregulated in dynein-2 mutants versus controls, shared between ciliated and non-ciliated cells**
